# High-level Prediction of Continuous Speech During Mind-Wandering

**DOI:** 10.1101/2025.08.29.673052

**Authors:** Gal R. Chen, Rachel Finkelstein, Ariel Goldstein, Ran R. Hassin, Leon Y. Deouell

## Abstract

Abundant evidence shows that when listening to speech or reading text, we continuously make predictions about upcoming words. Does this process stop when our mind wanders away and subjective experience becomes decoupled from the content of the speech? Conversely, is word prediction sufficient for conscious experience, or can these processes dissociate? Participants (N = 25) listened to audiobook content (> 12,000 words) while providing self-reports of mind-wandering. Mind-wandering was associated with spectral changes in the EEG signal, compared to attentive listening, as well as decreased early response to word onset, in line with current accounts. In contrast, EEG markers of word-level contextual surprise (indexed by the contextual surprise of large language models) remained intact during mind-wandering, alongside significant encoding of semantic content. Last, we measured the neural encoding of predictive context, measured by the correlation between EEG activity and the vectorial embeddings that language models use to predict each word given prior context. This encoding also persisted during mind-wandering, albeit weaker compared to attentive periods. Our findings show that the brain predicts and monitors relevant inputs even when the subjective experience of the external environment is dimmed during mind wandering. This suggests that the shared computational mechanisms between humans and Language Models are insufficient for having a subjective experience of the meaning of speech contents, with implications for theoretical accounts of mind-wandering and predictive processing accounts of consciousness.

**Significance Statement:** Large Language Models (LLMs) achieve human-like conversational abilities by using long range context to predict the most likely next word. Such predictive processing is also common in the brain, and some link the best prediction to the current conscious experience. Using EEG, we show that next-word prediction markers, marked by LLM-brain alignment, persist during periods of mind-wandering while people listen to audiobooks. Specifically, we found that when people report mind-wandering, their brains continue to represent context-based predictions, word-level surprisal, and word-level semantics. These findings dissociate high-level predictions from subjective experience.

## Introduction

It is nearly impossible to overestimate the importance of understanding what other people say, making the neural and cognitive underpinnings of speech processing a primary topic of interest for psychologists and cognitive neuroscientists (1–3). Importantly, speech processing involves not only the decoding of acoustic inputs, but also the conscious, subjective experience of what is being said (4, 5). Despite the progress made in understanding the basis of speech processing, it remains unclear which components of speech processing are associated with the subjective experience of engaged listening, and which can occur when consciousness is reduced. This question is especially relevant today, as the capabilities of deep learning models spark discussions regarding whether they can ultimately meet the criteria for consciousness (6–8).

Speech processing happens at multiple levels, including the physical properties of the stimulus (9), phonetics, word meaning (10, 11), and whole narratives (12). Here, we focus on the widespread use of contextual information (e.g., previous words) for extracting word meaning and predicting upcoming inputs (13, 14). Predictions are central to language comprehension (15–18), with contextual information influencing hierarchically lower levels of prediction errors (18). The theoretical and empirical importance of predictions raises a central question: Do high-level, context-based predictions co-occur with conscious experience? Or, alternatively, can we find instances where consciousness of speech content is attenuated, yet predictive and semantic processing persist?

Here, we use spontaneous fluctuations in attention to speech, known as mind-wandering (20), as a naturalistic case of reduced consciousness to relevant and audible external contents. The reader will likely recognize the experience of listening to a lecture or a podcast, only to realize at some point that they have no idea what was being said, having been absorbed in a stream of related or unrelated thoughts. Mind-wandering is usually defined as periods in which thought disengages from the current task (21), prompting descriptions of processing during mind-wandering as “perceptual decoupling,” in which perception becomes weaker, akin to ignored streams of information (22, 23). This description gains substantial support from studies showing decreased neural responsivity and behavioral performance during mind-wandering (24–27).

How deep is this perceptual decoupling? The literature on processing without awareness suggests that there is some evidence for high-level processing of singular objects (28–30), with integration being more difficult (30). However, such accounts are usually based on visual masking (28, 32) or concurrent external competition (33, 34), limiting the conclusions to rather weak signals (although, even under those conditions, there is recent evidence for high-level processing, e.g., 11, 35). Mind-wandering, on the other hand, includes reduced awareness of clearly perceptible contents.

Furthermore, mind-wandering reports are highly frequent during task performance (26, 36) and daily lives (37). This suggests that much can be gained if some form of high-level processing, such as next-word prediction and semantic processing (38, 39), persisted during mind-wandering. In the specific case of continuous speech, for example, monitoring goal- relevant content might allow for better allocation of resources during mind-wandering. It is possible, therefore, that the brain may still track and predict the meaning of single words, despite the reduction in subjective experience.

Here, we set out to examine the relationship between high-level predictive processing and mind-wandering in participants listening to audiobook chapters. We utilized previously established measures of pre-word context-related activity and post-word contextual surprise (19, 40) during self-reported periods of mind-wandering (MW) and On-task (OT). To quantify surprise, we utilized the probability of word occurrence assigned to each word by a Large Language Model (LLM, 41, 42). As a measure of semantic processing, we examine the neural encoding of word2vec (43) embeddings, which reflect semantic dimensions at the word level (44). Last, for context-related activity, we assessed the neural encoding of the representations that LLMs use to generate predictions, previously shown to share computational resources with humans (15, 45).

To foreshadow our results, we found that despite the decrease in response to word onset during MW, the other markers - pre-onset contextual activity, post-onset semantic encoding, and surprise processing - remained highly significant and even unchanged during MW. To the best of our knowledge, these findings are the first to show high-level verbal information processing, including prediction generation, during mind-wandering.

## Results

Our participants listened to 132 minutes of audiobooks, comprising six chapters, with occasional unexpected pauses during which they were asked to choose descriptions that best fit their experience at the moment of pause. Using validated methods for experience sampling (36), the choice “I was attentive to the audiobook”, was labeled as “on-task” (OT), while the rest of the options reflected different types of mind-wandering (all labeled MW, see Table S1). The MW-associated options were chosen in 28.1% of the time (SD = 10.0%), and the probability of MW responses was tightly correlated with post-listening ratings of interest at the audiobook level (r(4) = 0.96, Figure S1). Subjects had an average accuracy of 82.4% (SD = 7.4%) when examined about the audiobooks’ content (4-alternatives forced choice) after listening, indicating overall good compliance and understanding.

For all analyses below, we labeled the 12-second period prior to report as MW/OT based on participants’ response to the probe (for similar approaches, see review by Kam et al., 27). We begin by validating that MW reports reflect different states, as evidenced by changes in EEG spectra. Then, we move to examine semantic encoding, which is required for creating high-level predictions. The last two sections include an analysis of evoked responses to LLM-based surprisal at the word level (“surprise response” hereafter) and the encoding of context-related activity, both of which directly assess predictive processing during MW.

### Mind-wandering is Associated with Spectral Changes

We first tested whether the EEG signal includes any information regarding MW reports, supporting their reliability. Therefore, we examined whether mind-wandering reports can be predicted by the spectral properties of the EEG, as shown in previous studies (27, 46, 47). We split the Power Spectral Density (PSD) in each data segment (MW/OT segment) to familiar EEG bands (delta: 3-6Hz, theta: 6-8Hz, alpha: 8-12 Hz, low beta: 12-20Hz, and high beta: 20- 30 Hz, and the average broadband power between 3-30 Hz). A cluster-based spatial permutation analysis of the power difference between MW and OT (per band) revealed significant clusters of stronger power during the OT condition in the theta and low beta regime (Figure S2, lower panel). Beyond group differences, we were able to classify MW in all bands except for the delta and high beta bands in individual participants (all Bonferroni-corrected *p* < 0.01, see Figure S2, upper panel, and Table S3). While discussing the specific role of specific spectral bands in MW is outside the scope of this study, we note that both theta and beta bands are typically associated with sustained attention and executive control (48, 49).

Having validated that the EEG activity we recorded is sensitive enough to predict the subjective reports of our participants, we now proceed to examine word-level markers of semantic processing and contextual predictions during OT and MW periods.

### Semantic encoding persists during mind-wandering

A precondition for intact contextual processing is semantic processing of spoken words. To examine whether such processing occurs when participants report MW, we examined whether a model based on semantic properties of words can predict EEG responses left out of training. This was tested via an encoding model based on existing word2vec 100-dimensional embeddings, pre-trained on a Wikipedia corpus (50). We first extracted single-word epochs (from sections not labeled as MW/OT). Then, for each channel and timepoint (separately), we fitted a linear model predicting the EEG voltage with the semantic embeddings as features. We then tested this model on epochs labeled as MW or OT (outside of the training set), resulting in predicted voltage at each channel and timepoint. The average correlation between predicted and actual EEG, across channels, served as a marker for semantic encoding (Figure 2).

**Figure 1.**
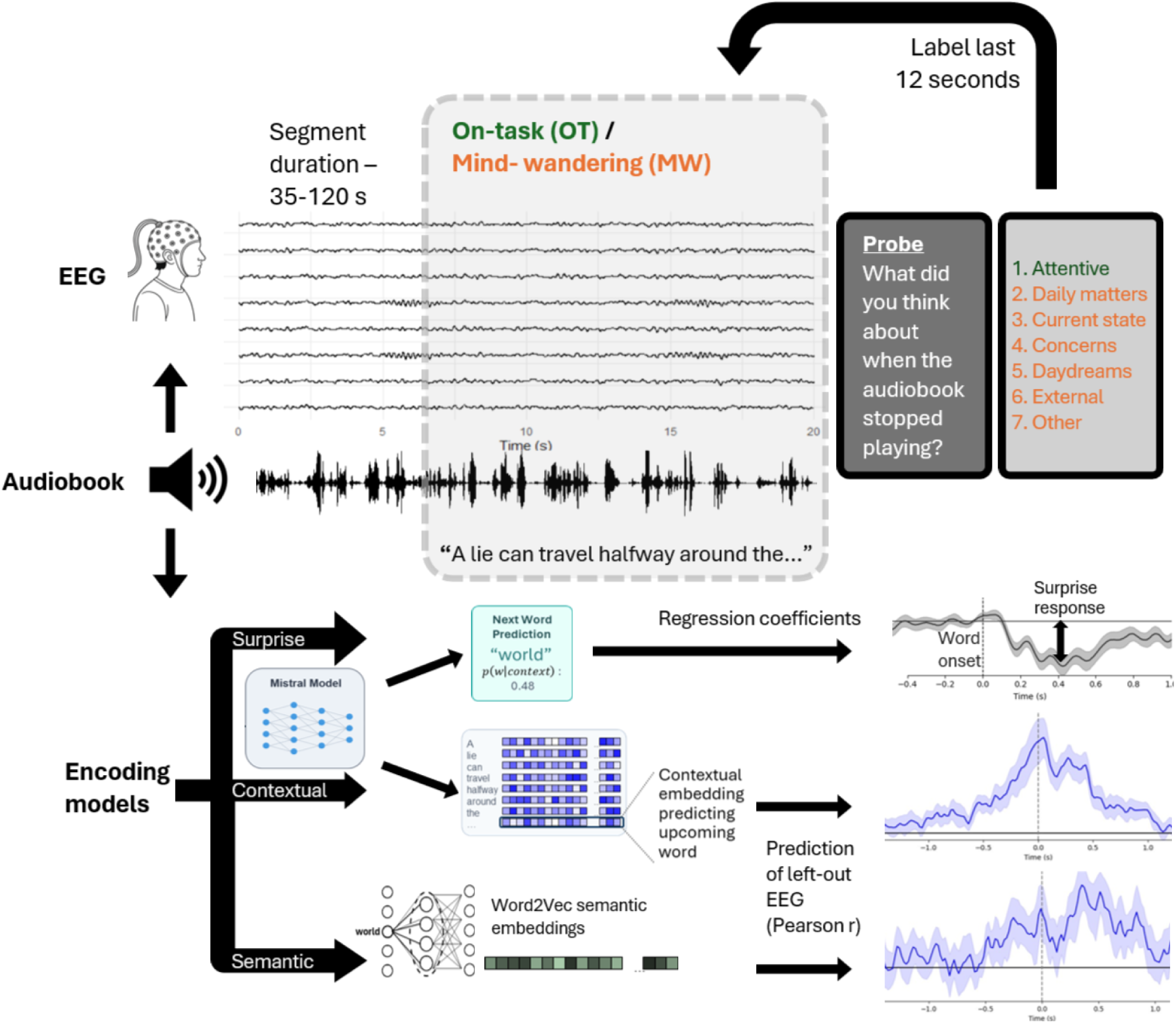
Experimental procedure and analysis approach. EEG was recorded from 25 subjects listening to six audiobook chapters, segmented into 136 parts varying in duration. The last 12 seconds of each segment were labeled according to a report in a post-segment probe. For each word played, we modeled three types of responses: change in response to surprising words (based on LLM), encoding of predictive context (based on LLM), and encoding of semantic embedding (based on Word2Vec). Each model was fitted separately. For surprise response, we focused on time-resolved coefficients estimating responses (separating MW/OT periods), and for semantic and contextual encoding, we tested left-out EEG data labeled as MW/OT (see Methods).

**Figure 2.**
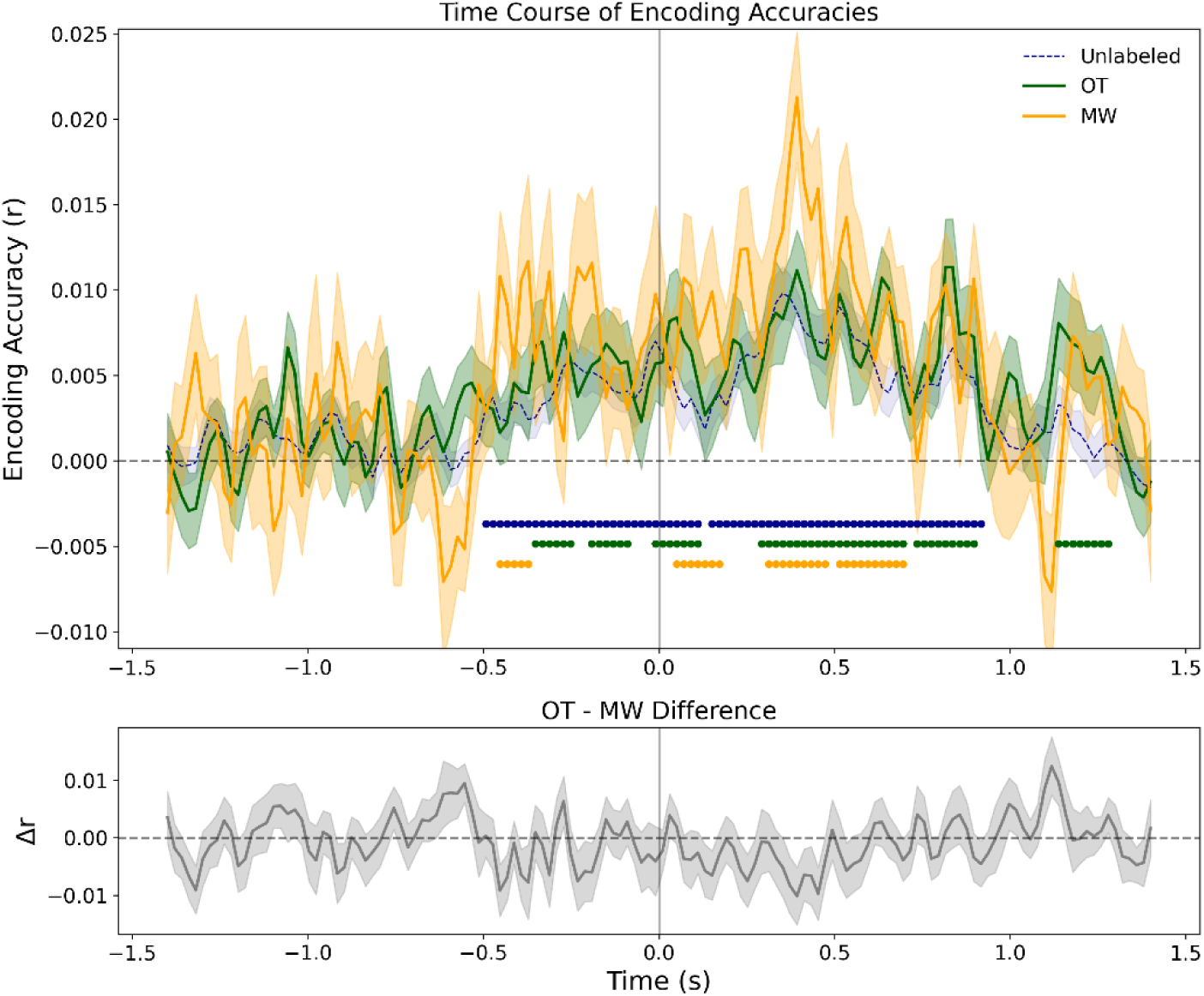
Semantic encoding persists during MW. Top: Average encoding scores (correlation between predicted and actual EEG) for word2vec semantic representations per time-point. Green and orange dots on the bottom denote the significance of the correlation, *p* < .05 after cluster-based temporal permutation test, for the OT and MW conditions, respectively. The results of the training data (blue, periods not marked as MW or OT), based on cross-validation, are shown for reference. Bottom: Difference between OT and MW correlations per timepoint. Shaded areas denote 1 standard error (SE) of the mean.

Similar to previous studies (15), encoding accuracy peaked at around 400 ms, associated with the time needed for lexical/semantic access (51). Semantic encoding was significant during both MW (significant clusters between 51-171 ms, 312-473 ms and 514-695 ms, all permutation *p* < 0.01, mean encoding between 300-500 ms, r = 0.0136, SD = 0.0118) and OT periods (significant clusters between 292-694 ms, permutation *p* < .001, mean encoding between 300-500 ms, r = 0.0083, SD = 0.009). Figure 2 denotes additional clusters with *p* < .05 in earlier timepoints.

Interestingly, peak semantic encoding during MW was not reduced compared to OT periods and, in fact, was numerically larger (temporal cluster-permutation, all *p* > .7; uncorrected difference between 300-500 ms: *t*(24) = -1.63, *p* = .11). Bayes factor analysis provided evidence against the hypothesis that OT encoding is stronger (BF10 = 0.089). Overall, this pattern of results is consistent with persistent semantic processing during MW.

### Mind-wandering reduces word onset response without affecting surprise response

So far, we have shown that semantic information is correlated with brain activity during MW, suggesting that this information may be available for generating predictions about upcoming words. Such predictions should be reflected in the evoked response to the words. We focus on two types of responses: first, the average response to a single word, capturing low-level processes that occur for all words. Second, the expected change in response when a particular word is more surprising. Both responses could be captured, respectively, by the intercept and slope of a simple regression model predicting EEG activity using surprise score (quantified by z-scoring the LLM-based contextual surprisal). To obtain these estimates, we conducted a time-resolved regression, estimating regression-based ERP (rERP), following a method tailored for overlapping responses (52), while also allowing for modeling the activity at each time point as a function of multiple variables. For each state (MW, OT and Unlabeled) the regression included predictors for word onset, separate predictors for surprise. We also included covariates for acoustic and lexical information and pre-word uncertainty (see Methods for additional details).

### Cluster-based analysis of onset response

To test the difference between onset responses while avoiding assumptions about time and location, the full waveforms in all channels were subjected to spatial-temporal permutation testing. Both MW and OT showed a significant response to word onset (MW: between 0-1000 ms, permutation *p <* .001; OT: between 0-949 ms, permutation *p <* .001), with a significant cluster for OT versus MW appearing during 51-379 ms (permutation *p* = .007, Figure 3a).

**Figure 3.**
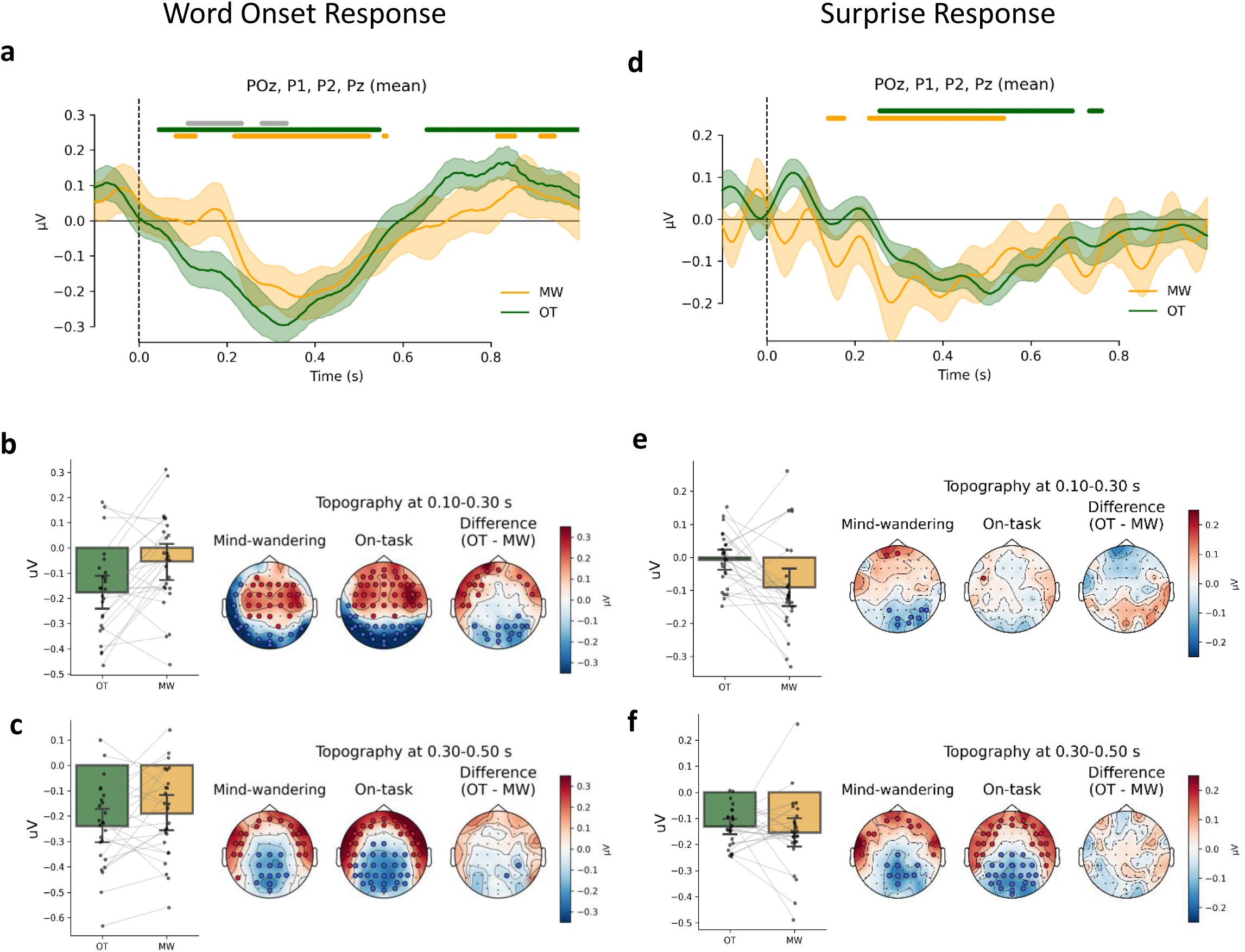
MW affects early response to words but not the response to the surprise level. (a) Grand average of rERPs for word onset predictor (binary) at posterior channels per attentional state. Time 0 reflects the onset of the word. Horizontal lines mark p < .05 at spatial-temporal cluster permutations for MW (orange), OT (green) and OT-MW difference (grey). (b) The effect of MW on the onset response. The bar graph shows the average estimated word onset response at the posterior cluster for the 100-300 ms time window for OT and MW words, with individual dots and lines reflecting single-subject estimates. The scalp topographies show the distribution of the rERP for each state and their differential activity. Red and blue dots mark channels belonging to a significant positive and negative clusters, respectively, during at least 50% of the time window (*p* < 0.05). (c) same as b, but for the 300-500 ms time window. (d-f) identical to a-c, but for the coefficients reflecting the surprise response, rather than the word onset response. All error bars and shaded areas in this figure denote 1 SE of the mean.

To gain further insight into the nature of these differences, we conducted an exploratory analysis using the average onset response at two different windows, which may help separate earlier processes from later, post-perceptual processes (100-300 and 300-500 ms, Figure 3b-c). We used the negative cluster found in the response to data unlabeled as MW and OT (excluding the last 12 seconds of each audiobook section), to select Pz, POz, P1, and P2 as the electrodes of interest for comparing the responses to the OT and MW conditions (see Figure S3 for other possible clusters of channels, and Figure S4 for the topographies for unlabeled data). At the early window, we found a stronger response for OT (mean OT-MW difference = 0.12 μV, *t*(24) = 3.23, *p* = .004, BF10 = 11.49). The effect of MW on onset response was numerically weaker also between 300-500 ms, but not significantly so (mean OT-MW difference = .049 μV, *t*(24) = 1.62, *p* = .117, BF10 = .67). We also tested the posterior cluster during the 200 ms before word onset with no apparent OT-MW difference (*t*(24) = 0.39, *p* = .69, BF10 = 0.23). Overall, in line with many studies showing attenuation of perceptual responses (27), MW is associated with a decline in early responses to spoken words.

### Cluster-based analysis of surprise response

To examine the response to word- surprisal during mind-wandering, we tested the coefficients of surprise in the regression model outlined above. We found an occipital-central negative coefficients associated with larger surprise at the word level (the coefficient reflects the estimated change in rERPs when a certain word is more surprising by one standard deviation). This effect was significant for both types of report (MW: between 242-761 ms, permutation *p* < .001; OT: between 152-789 ms, permutation *p* < .001, Figure 3d). The surprise response peaked somewhat earlier for MW than OT, based on the maximum of individual waveforms (mean latency difference = 70 ms, *t*(24) = 2.718, *p* = .012, BF10 = 4.10). No significant difference was found between OT and MW response amplitudes, suggesting no strong effect of MW on surprise processing (all *p* > .48). See Figure S5 for the effects of lexical frequency in the same model, which also showed no difference between OT and MW.

As for the word onset, we extracted the average difference at a posterior cluster of channels during two different time frames (100-300, 300-500, posterior cluster, Figure 3e-f, see Figure S6 for other clusters) and compared MW and OT responses. At the early window, mirroring the later peak observed for OT response, we found a stronger surprise response in MW (mean OT-MW difference between 100-300 ms = -0.084 μV [-0.15, -0.02], *t*(24) = -2.65, *p* = .014, BF10 = 3.61). Notably, this contrasts with the stronger onset response found for OT in the same time window. At the peak of the surprise response, between 300-500 ms, no difference was found, with a null effect more likely (mean OT-MW difference between 300-500 ms = -0.022 μV [-0.09, 0.05], *t*(24) = -0.67, *p* = .503, BF10 = 0.26). These results remained unchanged when function words were excluded and when controlling for words with high uncertainty, as quantified by the overall entropy of prior probability for word occurrence (Table S5). As with the onset response, we also tested the posterior cluster during the 200 ms before word onset with no apparent OT-MW difference (*t*(24) = 0.35, *p* = .72, BF10 = 0.22)

The above results indicate a significant decline in word onset response but no decline in the effect of surprise during mind wandering. To statistically test whether the MW effect differed significantly between the onset and surprise responses, we conducted a permutation test comparing the difference waves (MW effect on onset response minus MW effect on surprise response). This confirmed a larger effect of word onset than surprise level during the early time window (between 23-398 ms, permutation *p* = 0.007, Figure S7). Taken together, we found robust surprise-evoked negativity during both MW and OT periods, without a substantial decline during MW, despite the decrease in response to word onset.

### Analysis Using Variable Time Windows

One limitation of the results is that the 12- second window defined as MW may be contaminated by momentary OT periods. Contamination of this sort may create an apparent effect of surprise in our MW results, even if there is none in “pure” MW periods. This potential contamination should be less evident in shorter window durations, as subjects are more likely to accurately reflect on their experiences in the last few seconds prior to the probe. If contamination is responsible for our results, shortening the duration of the window should reduce the estimated effect of surprise during MW and/or increase the effect during OT periods.

To address this, we looked for an effect of pre-probe window on the surprise effect. For each probe, we assigned the subject’s choice of OT or MW to windows of 8-20 seconds pre-probe, in 2-second increments. We extracted the average amplitude of the surprise and onset responses at 300-500 ms after onset as the dependent variable. The effect of MW on word onset response was more substantial in the shorter time windows, consistent with it being less contaminated by OT data (slope of report x time-window interaction: *t*(148) = 2.50, *p* = .013; Figure 4), as indicated by a mixed-effect model predicting the amplitude by time window and report (MW/OT). For the surprise response, no such interaction was observed (*t*(148) = 0.21, *p* = .84). These results are inconsistent with a contamination explanation for the persistence of surprise effect during MW.

**Figure 4.**
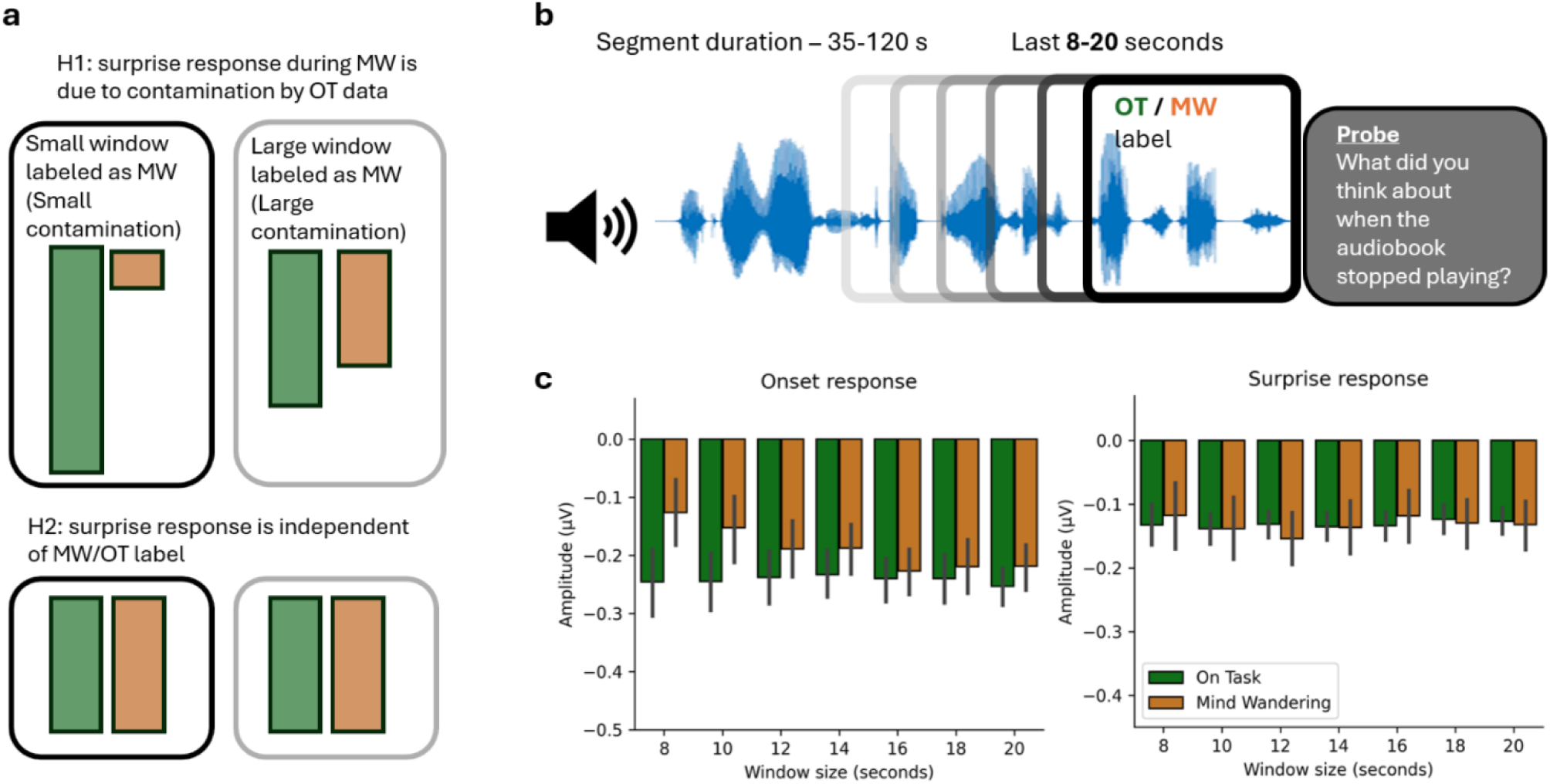
Surprise response during MW is independent of labeling window size. (a) Labeling a smaller temporal segment should result in less contamination of putative MW\OT periods by OT\WM periods. Under the hypothesis (H1) that the apparent effect of surprisal in MW periods is merely the result of contamination (with no genuine effect during MW), these effects should become smaller when the temporal window increases and larger when the window decreases. Conversely, if the surprisal effect persists during MW (H2), the size of the labeled segment should have little effect (b) Instead of the 12-second window used for the original analysis, we conducted the same analysis using windows of 8-20 seconds for labeling and extracted the mean amplitude of surprise and onset responses at the posterior cluster. (c) Analysis of MW effects per response and window. The y-axis shows the mean estimated response, and error bars denote one SE. Window size affects the onset response sensitivity to MW but not the surprise effect.

Overall, the analyses demonstrate that our design is sufficiently sensitive to capture the (somewhat expected) influences of mind-wandering on spectral power and evoked responses to speech. Despite this sensitivity, markers of word-level surprise persisted during MW, regardless of the time frame defined as MW.

### Mind-wandering attenuates but does not abolish contextual representations

The previous section established that MW does not decrease the sensitivity to word- level surprise, as reflected by the response to the words. Next, we asked whether the pre- word-onset contextual activity is also unaffected by mind-wandering. For this aim, we focused on the contextual information used by the LLM to generate the prediction, using a similar approach to the semantic encoding described above. If predictions are being conducted prior to word occurrence, then the features of these predictions should correlate with EEG activity to some degree. As contextual information, we used the last hidden layer of the Mistral (42, 53) model activations for each word, which includes the information used to predict the probabilities of upcoming words. These embeddings reflect just the context: they are identical for two different words with identical preceding contexts. Similar to the procedure used for semantic encoding, we used linear models trained on the embeddings of unlabeled EEG data (see Methods) to predict the EEG around single words and calculated the Pearson correlation between the estimated and actual EEG during MW and OT periods. This procedure was repeated for each timepoint and each channel, with high correlation meaning stronger neural encoding of embeddings, that is, stronger context-related activity.

This analysis yielded several insights. First, both MW and OT periods showed robust encoding of context, peaking around word onset (OT: -634 to 836 ms, permutation *p* < .001; MW: -372 to 71 and 131 to 574 ms, both with permutation *p* < .001, Figure 5). Second, while most timepoints did not show differences between OT and MW, our corrected cluster permutation test revealed a significant improvement in encoding for OT segments around the peak activity (between 30-151 ms, permutation *p* = .023). Specifically, the mean encoding around the peak, between -100 to 100 ms from word onset, was r = 0.0244 (SD = 0.0135) for OT periods, and r = 0.0162 (SD = 0.0181) for MW data (uncorrected difference, *t*(24) = -3.07, *p* = .005, BF10 = 8.4). Thus, we find that context-based predictive activity is somewhat diminished at word onset during mind-wandering, but still present.

**Figure 5.**
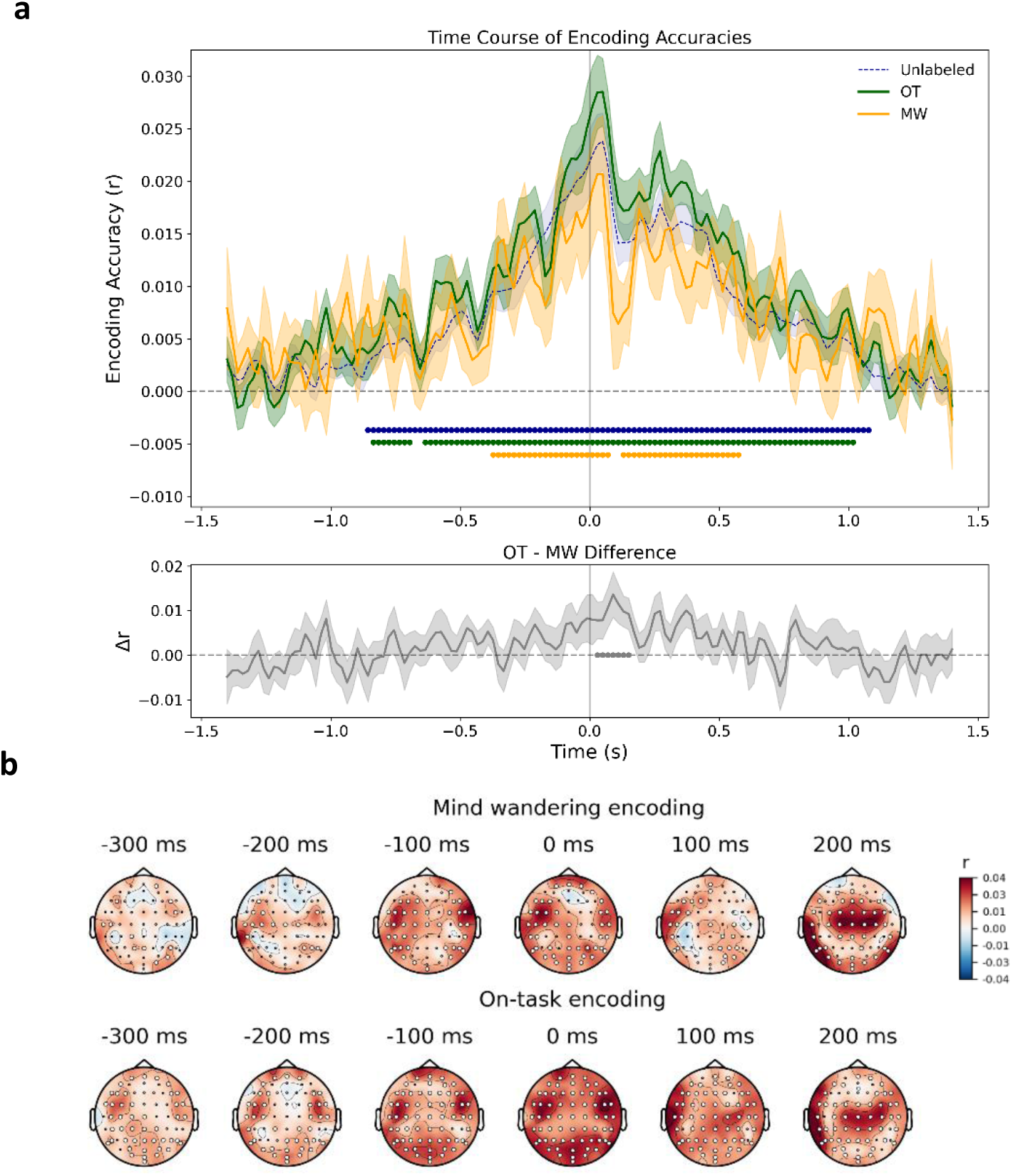
Encoding of contextual embeddings. (a) Results of average encoding scores per time-point. Green and orange dots denote *p* < .05 after cluster-based temporal permutation test. The grey line reflects the difference between OT and MW accuracy per timepoint, with black stars for significance. The results of the training data (blue, periods beyond the 12-second window we marked as MW or OT), based on cross-validation, are shown for reference. Shaded areas denote 1 SE. (b) shows the scalp topography of encoding scores in different timepoints for OT (bottom) and MW (top) periods, with white circles indicating *p* < .05 in spatial-temporal cluster-based permutation test.

We also conducted a control analysis with acoustic information of previous stimuli instead of contextual embeddings, as such information might be present in the contextual embeddings (54). We found that acoustic features performed worse in predicting EEG activity, compared to contextual embeddings, suggesting they could not account for all context-related activity (Figure S9).

## Discussion

Our results demonstrate that the brain continues to process continuous speech even when the mind wanders, and that it can use the extracted information to form predictions about the future. First, mind-wandering was associated with changes in the EEG spectrum and a weaker early response to words, likely reflecting perceptual decoupling. One might expect higher-level processing, involving word meaning, to become even weaker compared to early responses, which are associated with low-level processes. In contrast, we found that surprise response and semantic representation did not show a reduction, during mind-wandering. These results indicate that semantic and predictive processing do not necessarily indicate subjective experience

Not all processing markers we tested were unaffected by mind-wandering. Pre-onset contextual encoding was only partially preserved when participants’ minds wandered, compared to on-task periods, although it remained highly significant and followed a similar topography as in on-task periods. The difference was most substantial at word onset, possibly reflecting failure to align predictions with word onset, a reduction in prediction specificity, or, as a recent study suggested (35), a representation of a shorter context compared to attentive periods. Under these interpretations, mind-wandering impairs the ability to integrate recent and distant information to represent the changing context rather than single-word semantics.

The reduced contextual activation may seem at odds with the intact surprise response, which requires contextual prediction. This apparent discrepancy could be settled in two different ways. First, it is possible that the lack of a tight prior causes difficulty in lexical/semantic access for unpredicted information, compared to highly expected information. This difficulty may elicit more surprise-related activity during mind-wandering, as a previous study suggested (39). Second, it is possible that even weaker, but existing, contextual activity, is enough to generate surprise responses of the kind we observed. The exact link between pre-word contextual activity and surprise response is an important topic for further investigation.

### Perceptual Awareness and Processing Levels

The findings raise an important question about the depth of processing of external information during mind-wandering. The topography and timing of the surprise response were highly similar to the N400 ERP, associated with the difficulty of semantic integration or lexical access (11, 51, 55). The surprise response may also be associated with the prediction of low-level information based on context (e.g., predicting “Na” when “Napoleon” is expected), as a recent study (19) has demonstrated. Regardless of the level of processing of the word itself, the presence of a surprise response indicates that high-level information is being used to generate the prediction. When considered in conjunction with the persistent encoding of word2vec representations at around 400 ms, our results suggest that word meaning is indeed processed, at least to some degree, during mind-wandering.

When examined in the wider context of attention literature, our results seem at odds with studies showing robust decline in high-level speech processing (11, 56, 57), and even contextual predictions (35), when participants do not attend speech. It is important to note that, in contrast to many studies, we did not rely on selective attention or competition paradigms, which render the stimulus of interest task-irrelevant. Our findings, therefore, might serve as initial evidence that when participants do not actively suppress the information, processing might be more robust (58).

Our results carry immediate implications for decoupling accounts of mind-wandering (22, 24, 27), which implies that externally-oriented processes are inactive during such periods. We propose two possible hypotheses that explain how processing occurs during mind-wandering. One account suggests that while mind-wandering involves an internal stream of thoughts, we also remain aware of the external stream of information. This entails the possibility to simultaneously, or by rapidly alternating between contents, maintain awareness of two streams of information (56). Considering that we do not find a major effect of MW on predictive signals, this postulated mechanism appears to come without any processing cost. While split awareness in MW contradicts subjective intuition and participants’ reports, it warrants future scrutiny.

An alternative explanation holds that processing remains intact, despite diminished external awareness. This means high-level predictions are partially independent of conscious awareness (57), a view gaining support from studies of non-conscious speech processing (58–60). This interpretation may inform ongoing discussions about non-conscious processing capacity, which is mainly limited to the visual modality of singular events (61, 62). Further, it implies that theories describing high-level predictions as the basis for consciousness (63, 64) should be refined and address which types of predictions require or reflect conscious awareness.

The latter explanation makes our result particularly relevant to current discussions about AI consciousness (6, 7). LLMs are highly successful at verbal communication, promoting discussions regarding their ability to consciously experience and understand meaning (8). The capacity of such models stems from autoregressive predictions - that is, predicting the upcoming word given a broad context. Here, however, we show that in humans, such prediction capacities persist when individuals report not having a subjective experience of the text. This suggests that predictive capacities, on their own, cannot be used as an argument for LLMs having subjective experience, at least not in the same way humans do.

### Limitations

Our study has several limitations that future research might investigate. First, we address “prediction” as a single entity in this work, while it could be expanded to extract the specific semantic and linguistic dimensions being predicted (19). Because our focus is on *any* type of contextual prediction, we also did not separate between different parts of speech.

Second, we coded mind-wandering as a binary choice, to test if processing markers are diminished during *any* kind of mind-wandering. In practice, theoretical accounts often describe mind-wandering on multi-dimensional and continuous scales (65, 66). This introduces uncertainty regarding the depth of mind-wandering reports: in principle, some participants might wander only during the last few seconds, yet we labeled the previous 12 seconds as MW. We addressed this limitation by demonstrating that surprise processing remains robust even when using a shorter window, thereby making the periods more closely aligned with the report. Seeing that both time windows allowed for strong effects on the perceptual response, this contamination-based explanation cannot fully account for the strong processing markers found during MW. Still, MW reports cannot be argued as equal to reports of unawareness, highlighting the importance of assessing whether our results reflect purely non-conscious processing or processing during partial awareness. Future studies may also examine whether other measurement approaches of subjective state (e.g., continuous or multidimensional) can reveal differences within mind-wandering states.

We also note, with respect to the surprise response, that there are examples of a reduction in N400-like response during inattentive periods in general (51) and during audiobooks specifically (11, 67). Our study is distinguished from other works in two ways. First, we relied on spontaneous fluctuations in attention, rather than tasks requiring the selective suppression of “unattended” speech. Second, we conceptualized surprise as a continuous variable (avoiding artificial dichotomization) and controlled for overlapping words, which may have resulted in a more accurate estimation. However, other variables may also affect the results, such as the language used, engagement with content, and task duration.

## Conclusion

Our study provides evidence that multiple measures of semantic and contextual word processing remain robust during mind-wandering. As in many studies, reports of mind- wandering were indicative of reduced early responses and spectral changes in the EEG. Contextual prediction and semantic representation presumably reside higher in the processing hierarchy. Yet, we found that they remained intact, and possibly stronger, during mind- wandering. This suggests that while prediction is important for language processing, it is insufficient for the experience of being engaged with content.

## Materials and Methods

### Participants

25 participants took part in this experiment (17 females). All were native Hebrew speakers and students in the Hebrew University of Jerusalem and received payment (the equivalent of ∼$45) or course credit for participation. The study was approved by the Social Science Ethics Committee at the Hebrew University of Jerusalem, and all participants gave informed consent prior to participation. We chose a sample size of 25, which is larger than previous similar studies using audiobooks (11, 19).

### Stimuli and Procedure

Participants listened to six different audiobook chapters in the same order. The topics, by order, were: Urban legends, Greek drama (1^st^ chapter), Fiction writing, Product Economy, Greek drama (2^nd^ chapter) and Napoleon Bonaparte. All audiobooks were previously recorded by different native Hebrew speakers. The total duration of the heard chapters was two hours and 12 minutes (see Table S2). Listeners were instructed that they should do their best to stay attentive, and that they will be asked comprehension questions at the end.

All words in each audiobook were given a time-stamp by WhisperX (68). The timestamps and spellings were later checked and corrected by a human annotator, who corrected spelling mistakes and timing errors. For playing the sounds, we created stereo stimuli: one channel included the stimuli and was sent to the loudspeakers. The other channel included, at each starting point of words, impulse responses that were recorded in a separate channel of the EEG file. This allowed for precise timing of stimulus onsets.

Once every 40-120 seconds, with the exact distance sampled from an exponential distribution, the audiobook was stopped, and participants were asked about the contents of their experience in the moment of the pause. There were seven possible options, following an established measurement method (36). The first option was “I was attentive to the audiobook” and other six were “Daily matters”, “My current state”, “Personal concerns”, “Daydreams”, “External environment”, “Other” (Figure 1). The participants got a detailed description, read identically by the experimenter to ensure they understood that they should choose the first option whenever they felt attentive to the audiobook (See Table S1). We label the first option as “on-task” (OT), and all other options as “mind-wandering” (MW). We used the response to label the data in the 12-second window preceding the probe as either MW or OT. Time periods outside of the labeled window are heretofore described as “unlabeled data”. After the listening was over, participants answered 23 comprehension questions.

### Preprocessing and Analysis

EEG analysis was done using MNE-python version 1.4.0 (69). To pre-process the data, we first rejected noisy or completely silent channels by visual inspection. We then applied a 1- 49 Hz filter bandpass, interpolated rejected channels and applied an average reference. To clean activity associated with eye movements and blinks, we trained an Independent Component Analysis (70) on the raw data, and zeroed out the weights of components with topography and time course that reflect blinks or eye movement activity. The solution was applied to an unfiltered copy of the data. The clean data was then filtered between 0.5-8 Hz (see Table S4 for 0.5-20 Hz), the frequencies that were found to track speech responses (71, 72), which were also used in a recent prominent study (19) that examined contextual processing.

### Language Model Representations

For semantic encoding, we relied on word2vec embeddings, which capture context-independent semantic representations. For Hebrew word2vec embeddings, we used 100- dimensional vectors previously trained on co-occurrences in a Wikipedia corpus (50). To reduce the dimensionality for encoding, we used PCA on the training set (see below) to extract the top 40 components.

### Contextual embeddings

We extracted the latent representations (4096 dimensions) of a Hebrew version of Mistral 7B (42, 53) running on each audiobook with a window of 128 words in steps of one word. That is, for word n, the model was provided with words n-128 to n-1. The activations of the last hidden layer were taken for encoding analyses, as they provide the information the model uses for the prediction of word n.

### Word-level surprisal

Mistral 7B was also used to extract model-assigned probabilities of word occurrence, which were then transformed to negative log odds 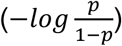 to create a contextual surprise score. The computation was based on a window of 25 words, which was found to saturate correlation with human performance in a next-word prediction task (15). The use of log odds means that the distance between probabilities of 0.01 and 0.1 is similar to the distance between 0.9 and 0.99, but larger than the distance between 0.45 and 0.54, which are qualitatively more similar (“somewhat certain”). To validate the surprise score, we conducted the same procedure on a HeBERT Model, extracting word-level surprisal. We found strong correlations in surprise scores given by the two models (all r>0.58 across audiobooks, Table S2). Word frequency was extracted using a corpus of frequencies in Wikipedia and movie subtitles (73), and was log-transformed.

### Spectrum-based Classification of Mind-wandering

To test for evidence that subjective mind-wandering reports were indeed associated with distinct patterns of brain activity, as previously demonstrated, we conducted a classification analysis within each subject based on the power in different channels and bands. For each subject and probe, we extracted data segments between 12 to 0 seconds before probe and assigned it a label based on the report. We then computed, per channel, the multitaper-based Power Spectral Density (PSD), and extracted the power in different frequencies and channels. For each frequency band, we extracted the mean logarithmic power in all channels. Our classification pipeline included a 10-fold cross-validation with 64 features (power in each channel) and 136 samples per subject. We used a Linear Discriminant Analysis (LDA) with Ledoit-Wolf shrinkage to train the classifier on 9 folds, and tested it on the left-out set, across all ten folds. LDA is especially suitable for our data because it uses prior probabilities for classes based on the training set proportions, accounting for imbalance. Our metric of success was based on the area under the curve of the receiver operating characteristic curve (ROC-AUC), a metric that is robust to classification in an unbalanced dataset.

### Null-hypothesis Significance Testing

The main statistical procedure in the paper was based on Maris & Oostenveld (74) cluster-based spatial-temporal permutation test when comparing the activity in two conditions within subject (hence, we analyzed difference waves). The procedure first locates and scores the sum of t-scores within a cluster, defined by points adjacent in time and/or space crossing the threshold of *p* = 0.05 in a two-tailed t-test. Then, the test performs a permutation-based procedure that randomly sign-flips the difference waves of individual subjects (effectively creating an expected difference of 0) and calculates the maximum cluster score in 5,000 iterations. For each of the original clusters, the corrected p-value represents the chances to get such a cluster by chance, that is, the percentile of the original result in the surrogate distribution created by the permutations. Whenever conducting single t-tests in follow-up tests, Bayes Factor is calculated based on a standard medium prior of 0.707(75).

### Encoding Analyses

Both the contextual and semantic encoding analyses were based on epoching of the data between -1400 to 1400 milliseconds relative to word onset (rejecting epochs with over 120 μV peak-to-peak differences) and fitting linear regressions for every timepoint and channel separately. Our cross-validation procedure used data from the beginning of each audiobook section (excluding the last 12 seconds), which was unlabeled as MW/OT, to train a model predicting the EEG of the test data (labeled as MW/OT) from embeddings. Specifically, we trained the regression model on the dimensionality-reduced representations (the first 40 PCA components) of the word embeddings in the data segments which were unlabeled as MW/OT (the whole dataset, excluding all segments of 12 seconds preceding probes). Then, we used the trained PCA weights to extract components representing the words in the test data (for MW and OT separately). The model based on unlabeled data was then used to predict the EEG activity of the labeled data, and Pearson correlation between the predicted and actual EEG was calculated as a metric of encoding success (at each timepoint and channel, across words). Importantly, the fact that the same model was used to predict MW and OT activity allows us to avoid potential confounds related to training set sizes.

For the main statistical analysis, since we had no prior hypothesis regarding the locus of the effect, we averaged the Pearson correlations across all channels as a representation of encoding accuracy, resulting in a singular time series of correlations per subject. A similar procedure was conducted using word2vec embeddings, which capture context-independent semantic representations. For Hebrew word2vec embeddings, we used 100-dimensional vectors previously trained on co-occurrences in a Wikipedia corpus (50). To reduce the dimensionality for encoding, we used PCA to extract the top 40 components. Like the contextual embeddings, PCA was trained only on the unlabeled (training) data to avoid leakage.

To account for the possible effect of autocorrelation in the EEG and the embeddings, we conducted a control analysis in which we shifted the words by a lag of 100 (so word n received the embedding of word n+100, Figure S8). This effectively breaks the relationship between the words and the embeddings.

For additional analysis of acoustic embeddings, we extracted the spectrogram of the previous word, which could evoke an activity that might be conflated with actual predictive context activity. Specifically, we used a Mel-spectrogram with 64 bands to extract acoustic embeddings. All spectrograms, for all words used, were reduced to 40 components using PCA. Then, a 6-fold cross-validation (each time leaving out one audiobook, fitting the PCA and linear model on the training data, and predicting the test data) procedure was applied to test the ability to predict the EEG activity from the embedding of the preceding word.

The main statistical procedure was a temporal cluster permutation test on the average time series, explained above, achieved by averaging all channels within subject. To gain further insight into the effects, we also calculated, for each subject, the average encoding score at 100 ms before and after the peak, that is, -100 to 100 for contextual and 300-500 for word2vec embeddings.

### Regression-based Analysis of Surprise response

To extract the response for contextual surprise, we conducted a multiple regression that accounts for overlap between words responses to consecutive words (52). It predicted the whole EEG time course with a design matrix that includes all word onsets as variables (coded as 1 in the design matrix), contextual surprise (continuous) as the primary variable of interest, with the log of lexical frequency also added to the model to control for its shared variance (∼10%) with surprisal. Additional covariates used were the time until the next word, average pitch (F0; using an autocorrelation-based algorithm (76), estimating pitch every 333 ms in steps of 111 ms), and two metrics of speech intensity. The speech intensity measures were mean intensity and sudden changes in intensity (via standard deviation of acoustic envelope), both of which were shown to explain the EEG signal (77). Pitch and both intensity metrics were calculated twice per word, once for the 500 ms preceding word onset, and once for the 500 ms after word onset, which resulted in 6 different predictors in the regressions. The regression modelled responses (and accounted for overlapping responses) from -500 to 1000 ms after word onset in each channel separately, rejecting segments with a peak-to-peak difference exceeding 120 μV.

Mind-wandering and OT periods were labeled based on a 12-second period before the probe. Data that fell outside these 12 seconds were unlabeled and utilized for training encoding models. For word onset, word frequency, and contextual surprise, three separate variables were used in the regression: one to model the response of unlabeled data, and one to model the reaction of MW or OT data. To allow differential effects, each variable was assigned a value of 0 when the occurring word belonged to one of the other labels. Frequency and contextual surprise were standardized (z-scored) before being split into the three MW conditions (MW, OT, Unlabeled). Due to the limited capacity of regressions to a high number of features, other covariates that were not of interest were not divided and used as one variable. The same splitting procedure was used to label data for the contextual embeddings and surprisal analyses. To ensure MW effects were not due to baseline differences occurring before word appearance, we calculated the mean activity between -200 and 0 ms in a posterior cluster of channels based on the unlabeled data topography (POz, P1, P2, and Pz, combined across channels).

## Supporting information

All supplemental tables

All supplemental figures

## Author Contributions

G.R.C., R.R.H., and L.Y.D. conceptualized and designed research,

G.R.C. and R.F. collected data and created software & materials, G.R.C. analyzed the data,

A.G. contributed new analytic tools, G.R.C., R.R.H., and L.Y.D. wrote the paper.

## Classification

Biological Sciences (Psychological and Cognitive Sciences)

