## Supplementary material for "High-level Prediction of Continuous Speech During Mind-Wandering": All supplemental tables

**Evidence from Neural Alignment with Language Models**

| Table S1. MW probes choice probability distribution | |
| --- | --- |
| **Option** | **Average Rate chosen** |
| 1 - Attentive to the chapter | 0.720 |
| 2 - Daily matters | 0.050 |
| 3 - My current state | 0.102 |
| 4 - Personal concerns | 0.019 |
| 5 - Daydreams | 0.064 |
| 6 - External environment | 0.029 |
| 7 - Other | 0.016 |

Optional choices in MW probes (instructions were only read in Hebrew)

**Hebrew:**

1. הייתי קשוב/ה לפרק – בכל מצב שבו העצירה הגיעה כשהיית מרוכז/ת בהאזנה, יש לבחור באופציה הזו.

2. דברים יום-יומיים – מחשבות על על סדר היום שלך, משימות שאת צריכה עוד לבצע היום בלי קשר לניסוי

3. מצבי הנוכחי – המצב המנטלי או הפיזי שלך (חם לי, קר לי, האלקטרודה מגרדת לי וכו')

4. דאגות אישיות – מחשבות מדאיגות על העתיד, ובכללי על החיים

5. חלומות בהקיץ – פנטזיות או מחשבות לא תלויות מציאות

6. הסביבה החיצונית – דברים שקשורים בחלל שאת נמצאת בו כרגע, ואובייקטים בו: השולחן, הרמקולים הקירות וכו'

7. אחר – כל מחשבה אחרת שלא נכללת באופציות הנ"ל ואינה ריכוז בתוכן הפרק הנוכחי

**English**

1. I was attentive to the chapter - **Choose this option whenever the pause occurred while you were focused on listening.**
2. Daily matters - Thoughts about your daily schedule, tasks you need to complete today unrelated to the experiment
3. My current state - Your mental or physical condition (I'm hot, I'm cold, the electrode is itching, etc.)
4. Personal concerns - Worrying thoughts about the future, and generally about life
5. Daydreams - Fantasies or thoughts unrelated to reality
6. External environment - Things related to the space you're currently in and objects within it: the table, speakers, walls, etc.
7. Other - Any other thought not included in the above options and not related to concentrating on the current chapter's content

| **Table S2. Details on each audiobook used.** | | | |  | |
| --- | --- | --- | --- | --- | --- |
| **Audiobook** | **Duration (seconds)** | **Number of words** | **Number of MW probes** | **Mean surprise /  negative log-odds probability (SD)** | **Correlation of Mistral-based surprise with heBert (Pearson’s r)** |
| Urban legends | 846.8 | 1476 | 19 | -1.06 (0.93) | 0.677 |
| Greek drama 1 | 1486.9 | 2816 | 28 | -1.07 (0.88) | 0.594 |
| Fiction writing | 1249.5 | 2631 | 23 | -1.02 (0.91) | 0.659 |
| Greek drama 2 | 1168.8 | 2347 | 22 | -1.15 (0.88) | 0.613 |
| Product economy | 1183.0 | 2284 | 22 | -0.99 (0.88) | 0.671 |
| Napoleon | 1995.2 | 2088 | 22 | -0.96 (0.85) | 0.585 |
| Total | 7930.2 | 13642 | 136 |  |  |

**Table S3. Results of within-subject MW classification based on PSD.**

| **Band** | **Mean AUC** | **Mean Sensitivity** | **Mean Specificity** | **t-score (AUC vs 0.5)** | **df** | **p** | **corrected p** |
| --- | --- | --- | --- | --- | --- | --- | --- |
| Delta  (3-6 Hz) | 0.53 | 0.651 | 0.946 | 1.287 | 24 | .2103 | 1 |
| Theta  (6-8 Hz) | 0.58 | 0.647 | 0.943 | 3.728 | 24 | .001 | .007 |
| Alpha  (8-12 Hz) | 0.596 | 0.62 | 0.949 | 5.385 | 24 | < .001 | < .001 |
| Beta1  (12-20 Hz) | 0.591 | 0.676 | 0.942 | 4.257 | 24 | < .001 | .0021 |
| Beta2  (20-30 Hz) | 0.547 | 0.611 | 0.942 | 1.992 | 24 | .0579 | .4053 |
| All  (3-30 Hz) | 0.58 | 0.624 | 0.941 | 3.823 | 24 | < .001 | .0056 |

| **Table S4.**  Comparison of main analysis using 0.5-8 Hz and 0.5-20 Hz filters. | | | | | | |
| --- | --- | --- | --- | --- | --- | --- |
| **Analysis type** | **MW/OT/ difference** | **0.5-8 Hz** | | **0.5-20 Hz** | | |
|  |  | **Cluster p-value** | **Timepoints (s)** | **Cluster  p-value** | **Timepoints (s)** | **Notes** |
| Word onset response | MW | < .001 | 0 – 0.95 | < .001 | 0 - 0.96 |  |
|  | OT | < .001 | 0 - 1 | < .001 | 0 - 1 |  |
|  | OT - MW | .007 | 0.05 - 0.37 | .003 | 0.06 - 0.37 | Direction: stronger early negativity for OT |
| Surprise response | MW | < .001 | 0.15 - 0.78 | < .001 | 0.23 - 0.60 |  |
|  | OT | < .001 | 0.24 - 0.76 | < .001 | 0.24 - 0.67 |  |
|  | OT - MW | 0.486 | 0.16 - 0.28 | .085 | 0.14 - 0.27 | Direction: stronger early negativity for MW |
| Contextual Encoding | MW | < .001 | -0.37 - 0.07 | < .001 | -0.39 - 0.05 | 1^st^ cluster |
|  | MW | < .001 | 0.13 - 0.49 | < .001 | 0.11 - 0.37 | 2^nd^ cluster |
|  | OT | < .001 | -0.63 - 0.83 | < .001 | -0.41 - 0.72 |  |
|  | OT - MW | .023 | 0.03 - 0.15 | .249 | 0.77 - 0.79 | Difference not replicated |
| Note. Reported are the clusters with maximum score from the cluster-based permutation analysis. The analysis is spatial-temporal for the rERP analyses, and temporal based on average of all channels for contextual encoding. | | | | | | |

| Table S5. Statistical Results for surprise response in different analyses | | | | | | | | |
| --- | --- | --- | --- | --- | --- | --- | --- | --- |
| **Time Window** | **Analysis Type** | **Mean OT** | **Mean MW** | **Mean  Difference** | **Cohen's d** | **BF₁₀** | **t(24)** | **p** |
| 100-300ms | Original | -0.007 | -0.091 | 0.083 | 0.713 | 3.607 | 2.651 | 0.014 |
|  | Only content words & high prior certainty words (z>-1) | -0.010 | -0.087 | 0.078 | 0.441 | 0.978 | 1.892 | 0.071 |
|  | Control for word class and certainty | -0.017 | -0.102 | 0.085 | 0.691 | 3.676 | 2.661 | 0.014 |
|  | Filter 0.5-20 Hz | -0.000 | -0.081 | 0.081 | 0.705 | 2.932 | 2.540 | 0.018 |
| 300-500ms | Original | -0.132 | -0.154 | 0.023 | 0.193 | 0.260 | 0.680 | 0.503 |
|  | Only content words & high prior certainty words (z>-1) | -0.125 | -0.153 | 0.028 | 0.142 | 0.236 | 0.493 | 0.626 |
|  | Control for word class and certainty | -0.125 | -0.149 | 0.023 | 0.191 | 0.264 | 0.703 | 0.489 |
|  | Filter 0.5-20 Hz | -0.122 | -0.148 | 0.025 | 0.214 | 0.274 | 0.758 | 0.456 |

| Table S6. Statistical Results for word onset response in different analyses | | | | | | | | | |
| --- | --- | --- | --- | --- | --- | --- | --- | --- | --- |
| **Time Window** | **Analysis Type** | **Mean OT** | **Mean MW** | **Mean  Difference MW-OT** | **Cohen's d** | **BF₁₀** | **t(24)** | | **p** |
| 100-300ms | Original | -0.177 | -0.053 | -0.124 | 0.691 | 11.488 | -3.230 | | 0.004 |
|  | Only content words & high prior certainty words (z>-1) | -0.123 | -0.016 | -0.108 | 0.556 | 1.827 | -2.276 | | 0.032 |
|  | Control for word class and certainty | -0.179 | -0.056 | -0.123 | 0.686 | 10.524 | -3.188 | | 0.004 |
|  | Filter 0.5-20 Hz | -0.180 | -0.052 | -0.128 | 0.703 | 12.895 | -3.285 | | 0.003 |
| 300-500ms | Original | -0.238 | -0.189 | -0.049 | 0.287 | 0.667 | -1.625 | | 0.117 |
|  | Only content words & high prior certainty words (z>-1) | -0.150 | -0.102 | -0.048 | 0.282 | 0.437 | -1.281 | | 0.212 |
|  | Control for word class and certainty | -0.239 | -0.191 | -0.049 | 0.286 | 0.660 | -1.617 | | 0.119 |
|  | Filter 0.5-20 Hz | -0.242 | -0.190 | -0.053 | 0.306 | 0.795 | -1.751 | | 0.093 |
