## Supplementary material for "High-level Prediction of Continuous Speech During Mind-Wandering": All supplemental figures

**Gal R. Chen^1^, Rachel Finkelstein^2^, Ariel Goldstein^3^, Ran R. Hassin^1,4^, Leon Y. Deouell^1,2^**
^1^ Department of Psychology, The Hebrew University of Jerusalem

^2^ Edmond and Lily Safra Center for Brain Sciences, The Hebrew University of Jerusalem

^3^ Department of Cognitive and Brain Sciences and Business School, The Hebrew University of Jerusalem

^4^ The Federmann Center for the Study of Rationality, The Hebrew University of Jerusalem

*Corresponding author: Gal R. Chen

**Figure S1. Behavioral Analysis of MW reports**

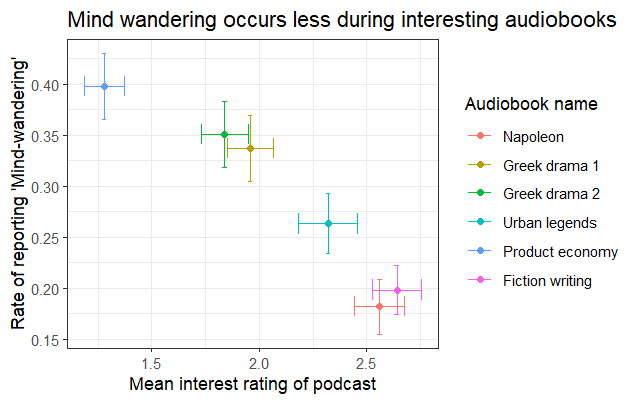

At the end of the Experiment, subjects were presented with the title of each chapter, alongside a short reminder of the topic, and were asked to rate how engaging the content of this audiobook chapter was. Options were 1 (Not interesting at all), 2 (Somewhat interesting), or 3 (Very interesting). We averaged the scores for each audiobook, and did the same for the rate of MW reported in each audiobook. The Pearson correlation between the two measures was r = 0.96, *t*(4) = 7.2, *p* = 0.002. Each point in the graph is one audiobook, and the error bars reflect standard error across subjects. The result indicates that more interesting audiobooks were also ones in which subjects were less likely to mind-wander.

**Figure S2. Mind-wandering classification**

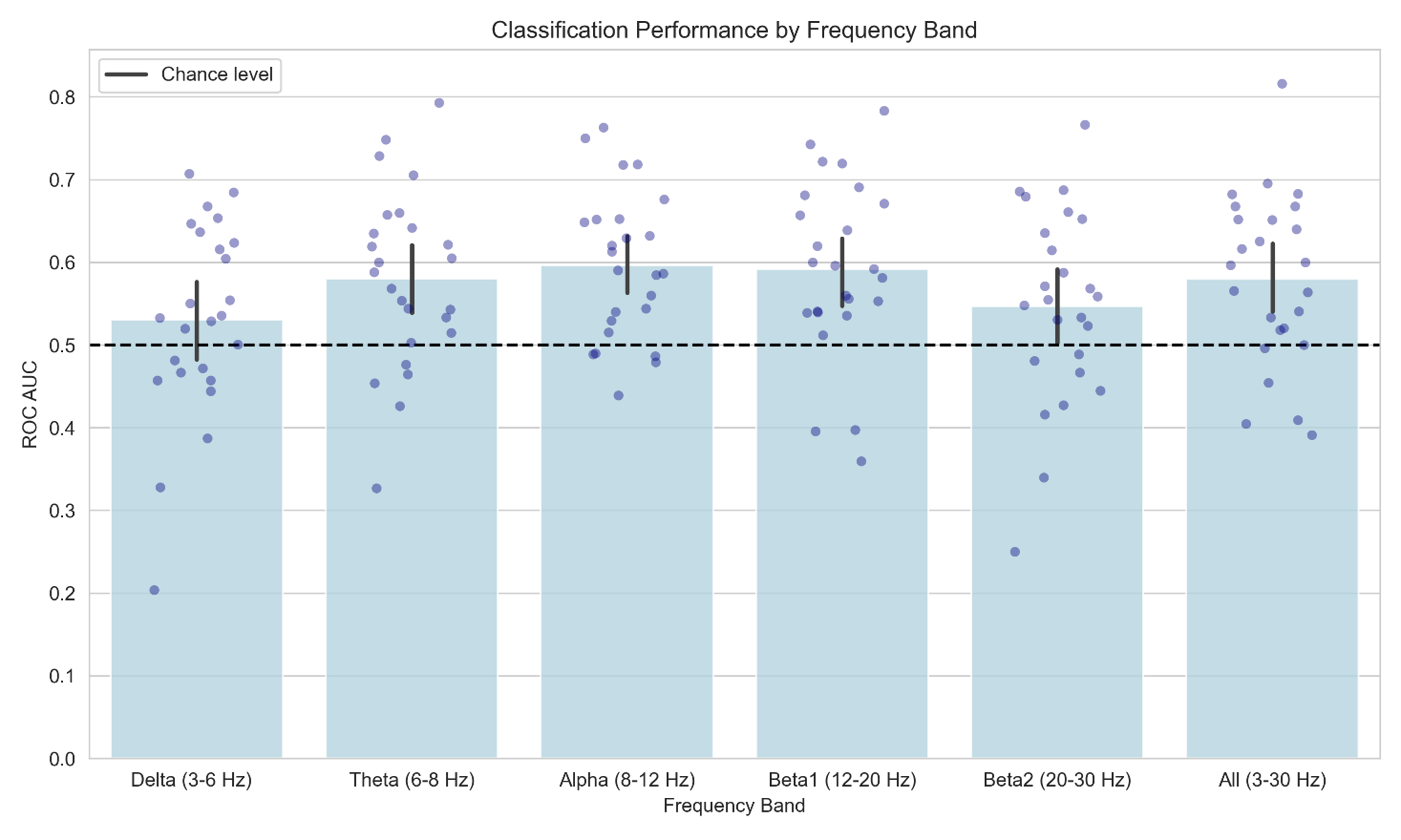

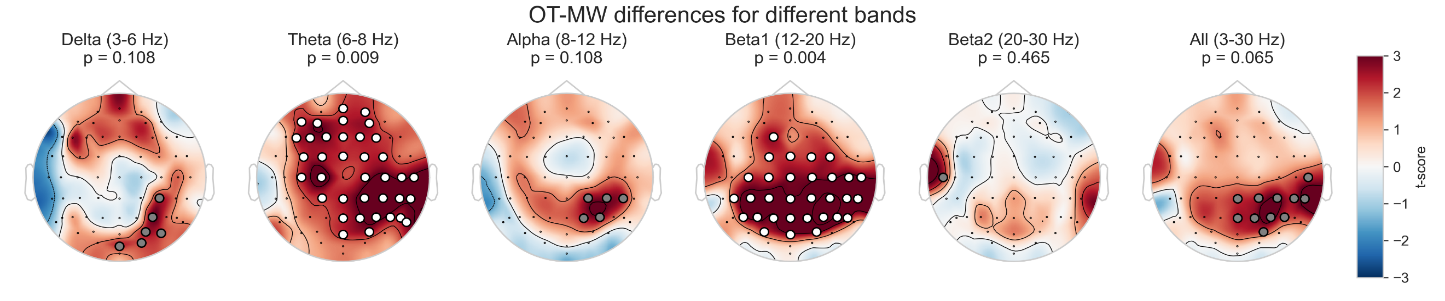

Top: Accuracy (ROC-AUC) of single-epoch (12 seconds) classification based on the average power in the epochs. The classification was based on a 10-fold cross-validation, with the average power in each band across all 64 channels. Error bars denote 95% confidence interval. Bottom panel shows the average difference in power between OT and MW reports, split between different bands. White circles denote channels belong to a significant (corrected *p* < .05) cluster after spatial cluster-based testing. Grey circles show the maximum cluster (uncorrected *p* < .05) for non-significant tests.

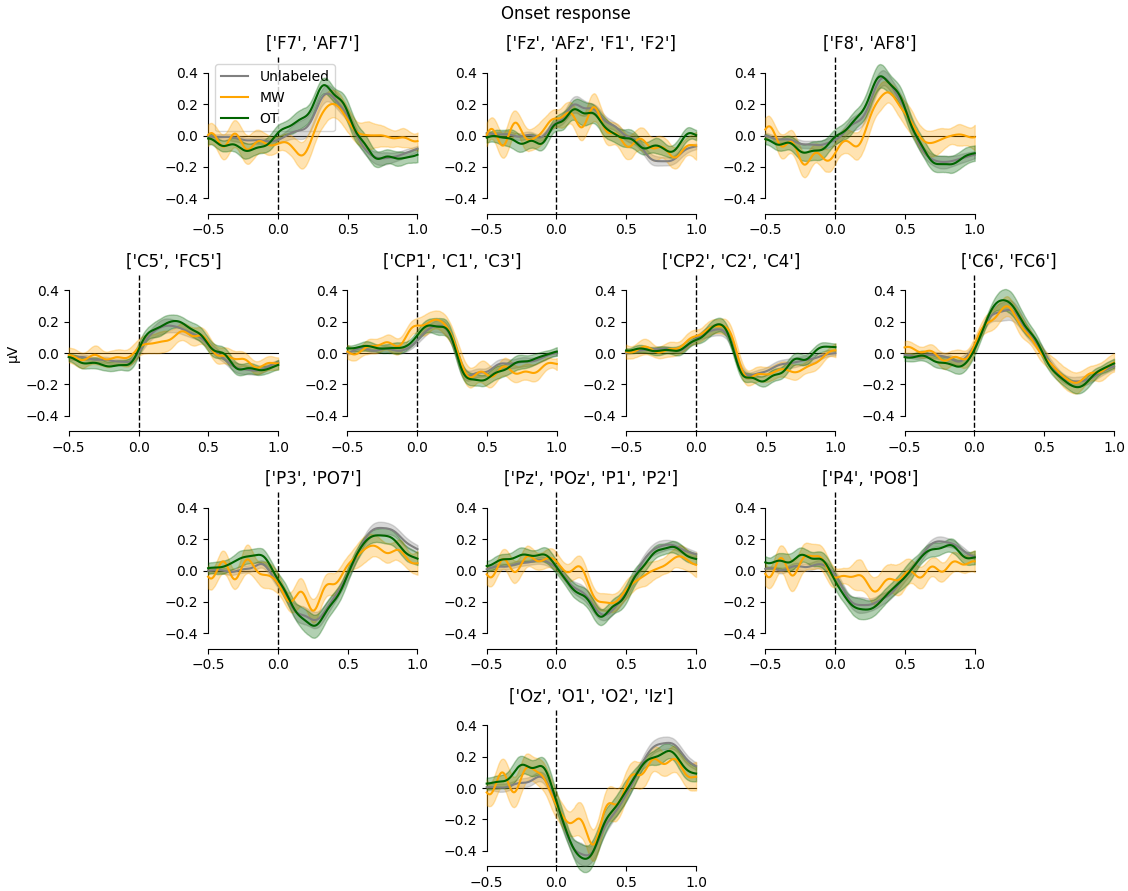
**Figure S3. Responses to word onset in different clusters of spatially adjacent channels.**

This figure is identical to Figure 4a, but each individual plots is based on different cluster of channels.

**Figure S4. Topography of surprise and onset responses for unlabeled data.**

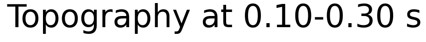

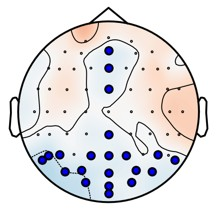

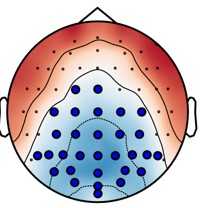

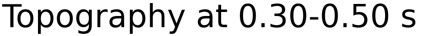

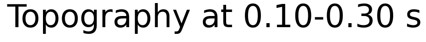

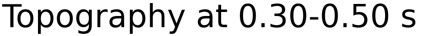

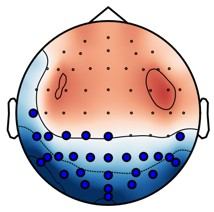

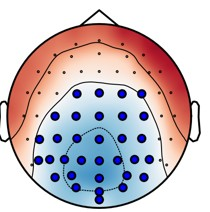

Surprise response

Onset response

Note. The marked dots are based on a similar spatio-temporal cluster permutation test conducted for MW and OT data in Figure 4. They are based on coefficients for which the design matrix zeroed for words belonging to MW/OT periods, and so they reflect another portion of the dataset.

**Figure S5. Lexical frequency analysis**

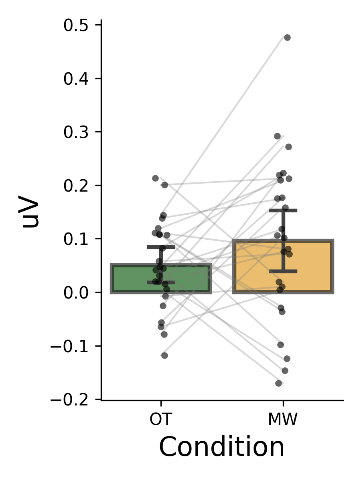

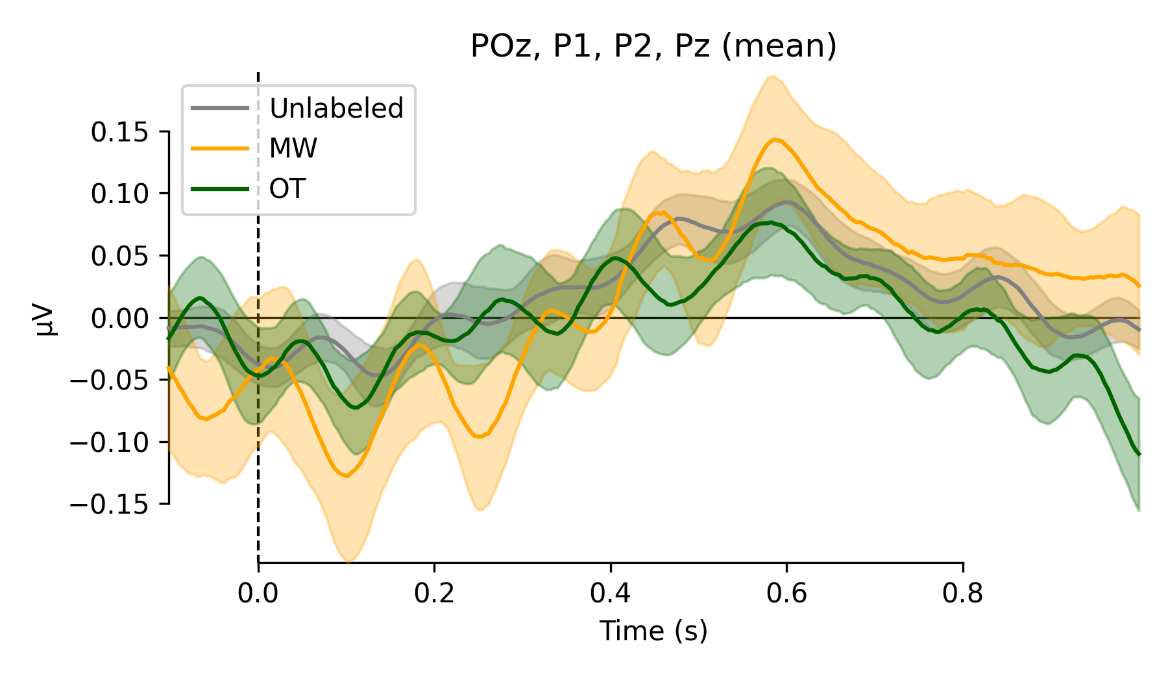

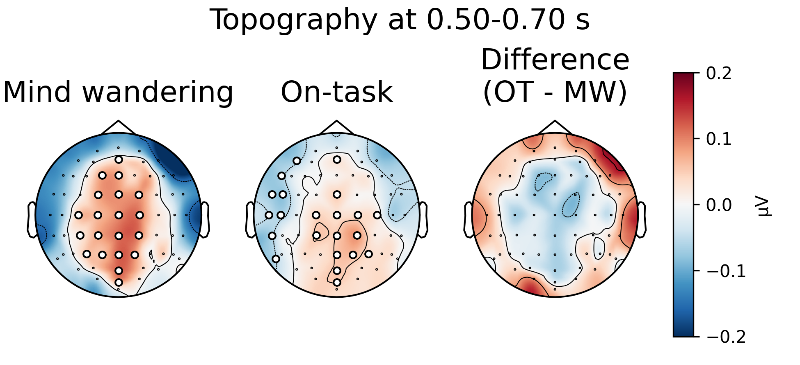

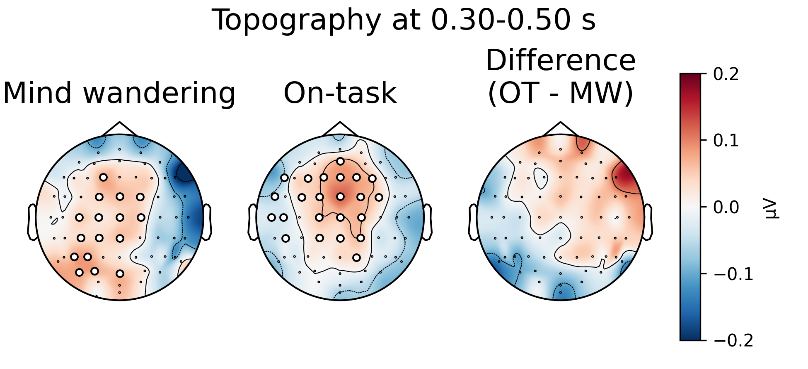

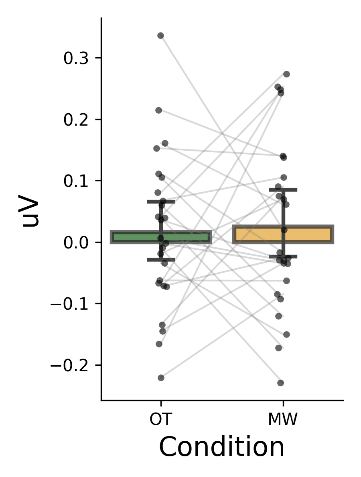

Note. The figure presents the effect of lexical frequency, included in the model with surprise and other reported covariates, on the EEG response to words. It is identical to Figure 4 in its format. No significant differences were found in spatial-temporal cluster permutation test.

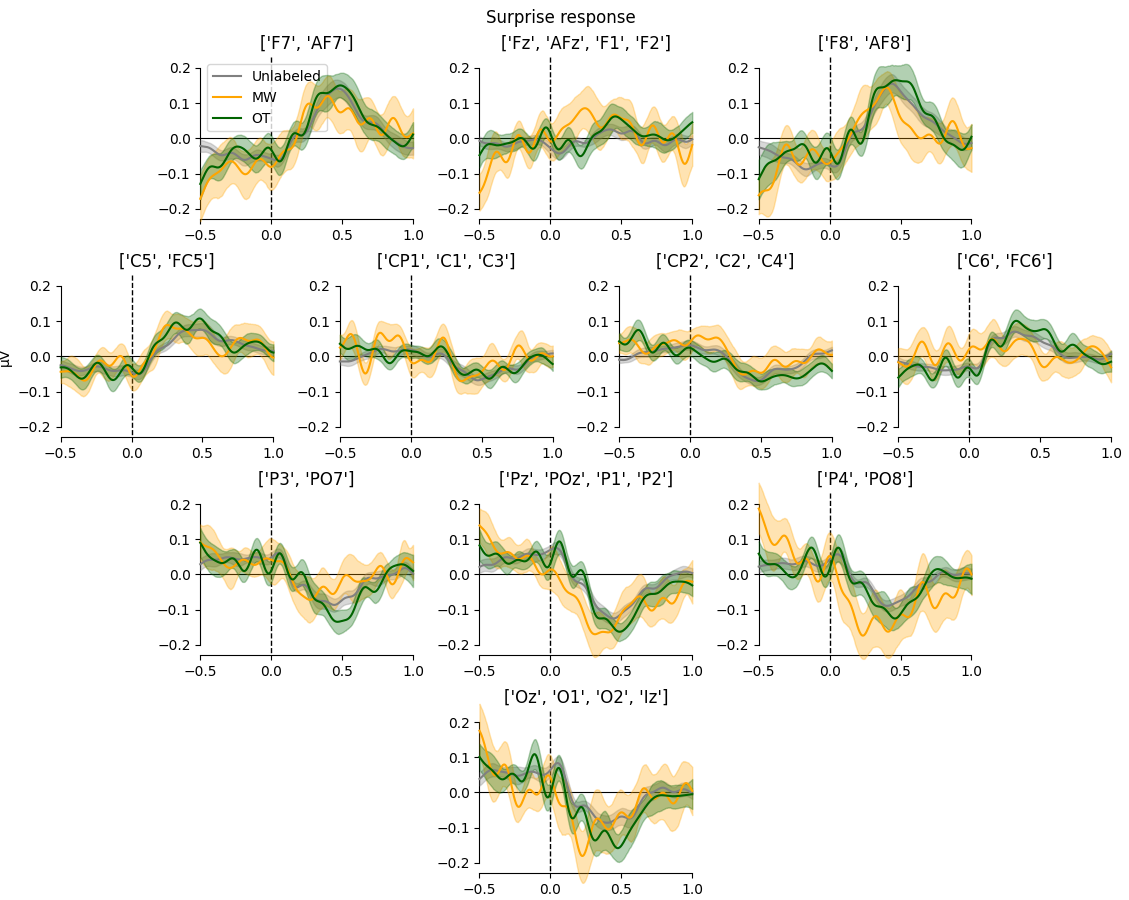
**Figure S6. Responses to contextual surprise in different clusters of spatially adjacent channels.**

This figure is identical to Figure 4b, but each individual plots is based on different cluster of channels.

**Figure S7. The difference between MW effect on word onset and MW effect on contextual surprise.**

Note. The top panel shows the average difference waves, based OT-MW difference for surprise (orange) and onset (blue) response. The green line shows the difference between surprise and onset effects (the interaction effect). The bottom panel shows the interaction difference wave topography, with the significant clusters in spatial-temporal cluster permutation test (*p* < 0.05) appearing in white dots.

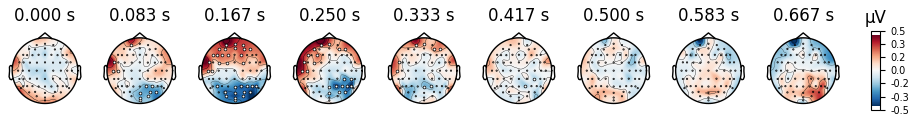

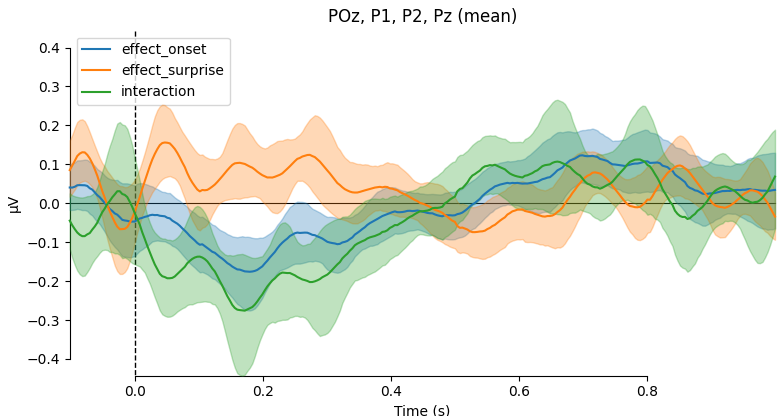

**Figure S8. Encoding of lagged contextual embeddings.**

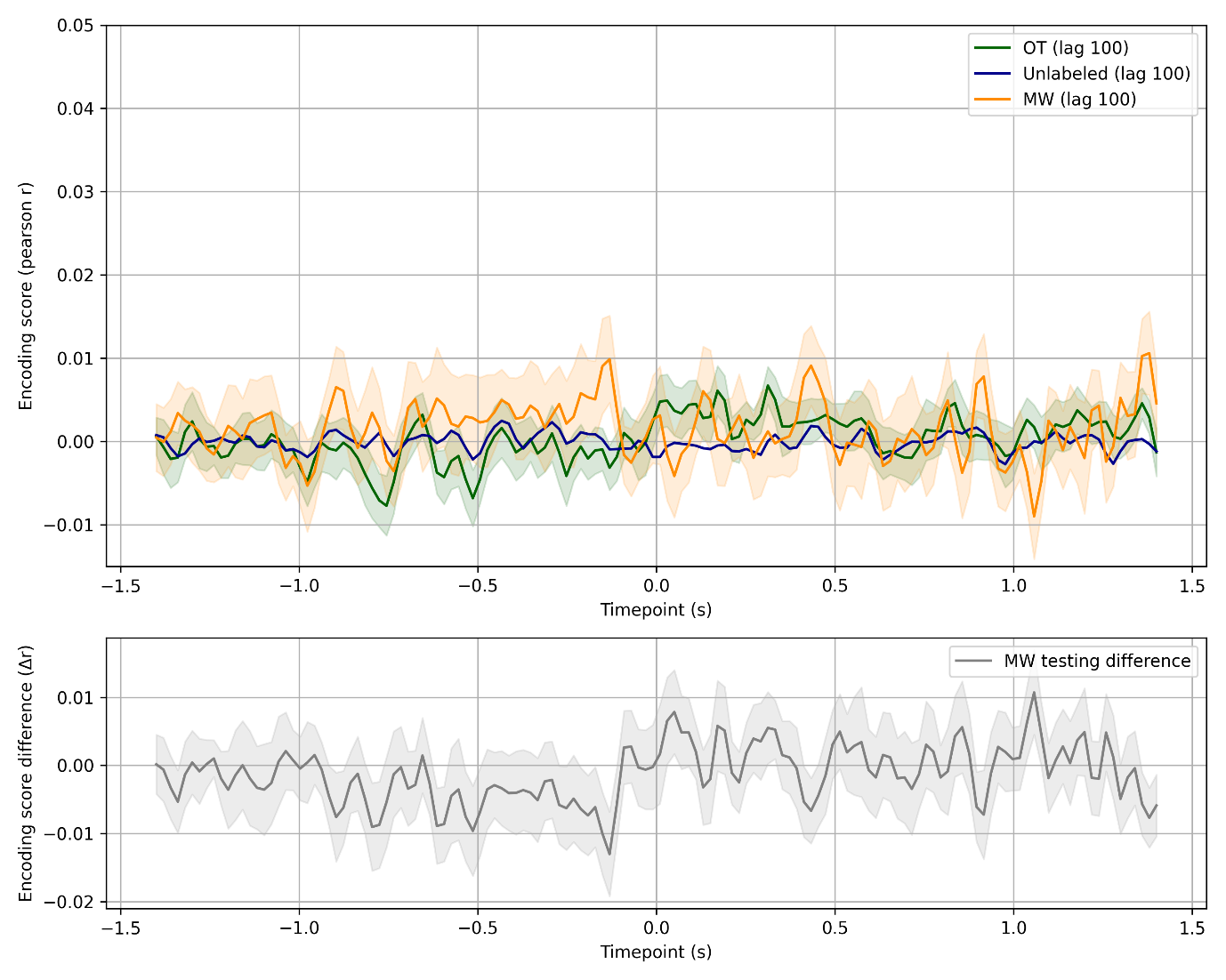

Note. This analysis is identical to the one used in Figure 5 (contextual embedding analysis), but instead of using the embeddings of the word n for analyzing epoch n, we used the embedding of word n+100. As such, the last 100 word in each audiobook are discarded from this analysis.

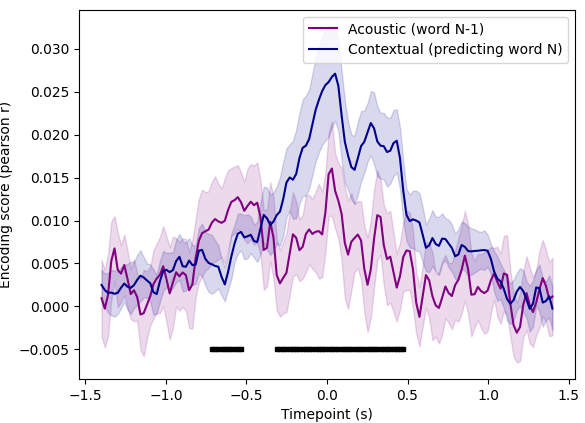
**Figure S9. Encoding of acoustic embeddings.**

Note. This analysis is based on the first 40 principal components of mel-spectrogram at 0-350 ms post word onset, for all data without regards to MW label. It shows that the acoustic information of the previous word cannot account for the time course of contextual encoding. Black squares denote timepoints in which the difference is significant at cluster permutation testing. Both accuracies are based on 6-fold cross validation.
